# Atomic force microscopy reveals the mechanical properties of breast cancer bone metastases

**DOI:** 10.1101/2021.06.01.446539

**Authors:** Xinyue Chen, Russell Hughes, Nic Mullin, Rhoda J. Hawkins, Ingunn Holen, Nicola J. Brown, Jamie K. Hobbs

## Abstract

Mechanically dependent processes are essential in cancer metastases. However, reliable mechanical characterisation of metastatic cancer remains challenging whilst maintaining the tissue complexity and an intact sample. Using atomic force microscopy, we quantified the micro-mechanical properties of relatively intact metastatic breast tumours and their surrounding bone microenvironment isolated from mice, and compared with other breast cancer models both *ex vivo* and *in vitro*. A unique mechanical distribution of extremely low elastic modulus and viscosity was identified on metastatic tumours, which were significantly more compliant than both 2D *in vitro* cultured cancer cells and subcutaneous tumour explants. The presence of mechanically distinct metastatic tumour did not result in alterations of the mechanical properties of the surrounding microenvironment at meso-scale distances (> 200 µm). These findings demonstrate the utility of atomic force microscopy in studies of complex tissues and provide new insights into the mechanical properties of cancer metastases in bone.

## INTRODUCTION

Mechanics are widely agreed to play a critical role in cancer development, progression and metastasis^1^. Existing studies of cancer mechanics commonly measure either the micro-properties of single cultured cells, or the bulk properties of whole tumour tissues. Single cultured cancer cells have repeatedly been shown to be more compliant than their healthy counterparts^2, 3, 4, 5, 6^, whilst tumours (*in vivo*/*ex vivo*) are stiffer than their surrounding tissues^1, 7, 8^. This paradox has commonly been explained by tumours exhibiting stiff components (e.g. collagen) in the peripheral stroma^1, 9^. A comprehensive mechanical study of primary breast tumours, using relatively intact human biopsies, reported that cancer progression is associated with a significant softening of tumour epithelial cells^10^, but is not limited to the previously assumed matrix stiffening^9^. This demonstrates the heterogeneity of the mechanical properties within the tumour microenvironment, and indicates the importance of bridging across length scales when performing mechanical measurements.

To improve our understanding of the role that mechanics play in the later stages of cancer development, reliable mechanical characterisation of metastatic tumours and the surrounding microenvironment is essential, but currently there are few published studies. Bone is one of the most common metastatic sites for multiple cancers, including breast cancer^11, 12^, with metastasis increasing patient morbidity and mortality. Once cancer has spread to the skeleton, it is considered incurable, so methods to characterise this process which allow the identification of effective therapeutic interventions need to be developed. Due to the complex nature of the multicellular bone microenvironment, interactions between tumour and bone cells have been widely explored as key to driving metastatic growth and therefore represent therapeutic targets (recently reviewed by Coleman *et al*^13^). As it is impossible to carry out detailed analyses of tumour spread to bone in humans, murine model systems mimicking the different stages of bone metastasis have been developed^14^. We and others have used these models to demonstrate how breast cancer cells colonise specific areas of bone (niches) that support their survival and progression, ultimately resulting in tumour-induced bone destruction in the form of lytic lesions that can be visualised using µCT analysis^15, 16, 17^. It is extremely challenging to access bone metastases without perturbation, due to the complex structure of bone (i.e. the hard shell surrounding the near liquid bone marrow). Meanwhile, where mechanical characterisation on complex tumours has been performed as in the study by Plodinec *et al*^10^, the viscoelastic effects that occur during tumour progression^18, 19^ were not taken into account.

Here, we have quantified the micro-mechanical properties, including both elastic moduli and viscosity, of relatively intact breast cancer experimental bone metastases and the surrounding microenvironment in a mouse model^15, 20^ using colloidal probe atomic force microscopy (AFM)^21^. The results were compared to explanted subcutaneous breast tumours (grown in mice) and single 2D cultured breast cancer cells in a petri-dish, revealing significant differences in mechanical properties between simplified 2D *in vitro* models, non-orthotopic tumours (i.e. breast tumour grown under the skin) and metastatic tumours (both 3D). We also revealed, by comparison to the properties of normal tumour-free bone microenvironment, that in our model system there is minimal mechanical impact of the metastatic tumour on its surrounding environment at meso-scale distances (> 200 µm).

## RESULTS

### AFM reveals extremely low elastic modulus and viscosity in bone metastases

Breast cancer bone metastases were established in a commonly used *in vivo* mouse model^15, 20^ (Fig. 1a-b). As illustrated in figure 1c-g, metastatic tumour foci develop in close proximity to trabecular bone structures in the metaphysis region of the long bones (tibia and femur) 3-4 weeks after intra-cardiac injection of MDA-MB-231^luc/GFP^ breast cancer cells. To probe the mechanical profile of the metastatic tumour (MT), we applied point force (*F*) *vs*. indentation (*δ*) and creep measurements by AFM, as described in our previous study characterising the micro-mechanical properties of tumour-free bones^21^. This is a robust method for measuring mechanical properties of complex tissue, provided sufficient biological and technical repeats are utilised. AFM measurements were applied to the surfaces of split fresh bones *ex-vivo*, at randomly selected positions (in total 126 positions from 19 bone samples) within the tumour area (Fig. 1f-g). The Young’s modulus *E*_*H-S*_ was obtained from Hertz-Sneddon (H-S) model fits to the *F*-*δ* curves. The viscoelastic properties, including the Young’s modulus *E*_*K-V*_ and viscosity *η*, were obtained from Kelvin-Voigt (K-V) model fits to the indentation change (*Δδ*) *vs*. creep time (*t*) curves.

**Figure 1.**
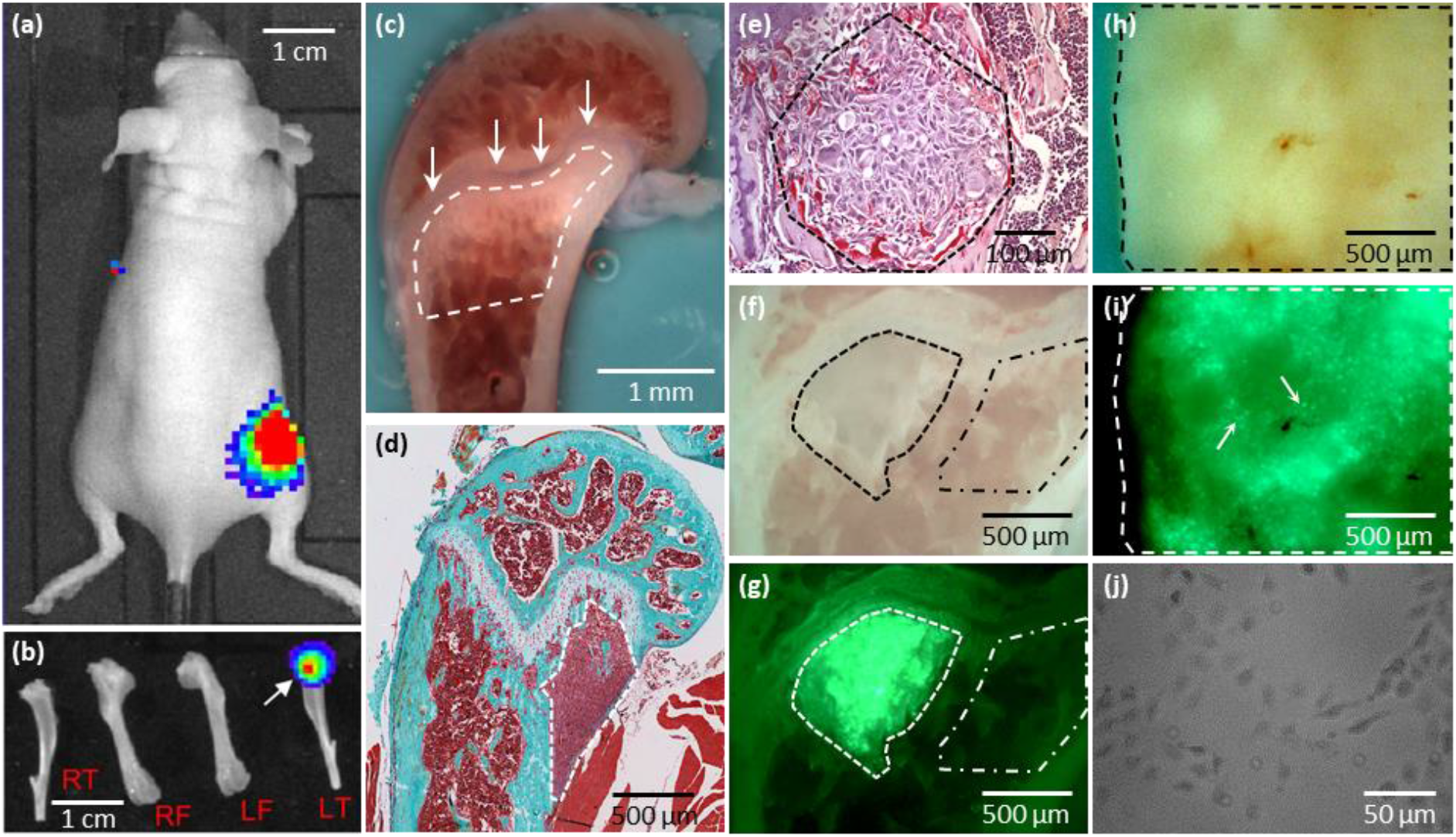
Illustrative optical and histological images of the different cancer models used in this study. (a) *In vivo* bioluminescence image of a mouse with breast cancer metastasis to bone (in *colours*). (b) *Ex vivo* bioluminescence image of the dissected hind limbs of the mouse in (a) (RF: right femur; RT: right tibia; LF: left femur; LT: left tibia). The bioluminescent signal (in *colours, arrow*) indicates the LT contains breast cancer metastases at the proximal end. (c) An example optical image of the bone surface (femur) from a tumour-free mouse *in situ* on the AFM. The region of interest in this study is the metaphysis (outlined by *dashed line*) located just below the growth plate (*arrows*), which is predominantly colonised by metastatic cancer cells. Examples of (d) Goldner’s trichrome stained histological section and (e) tartrate-resistant acid phosphatase stained higher magnification histological image of breast cancer metastases in bones. The metastatic tumours (MT) are outlined as dashed regions. Images (d-e) adapted by permission from I. Holen: Springer Nature Ref ^33^, Copyright 2012. (f) Conventional and (g) corresponding *in situ* fluorescent images of an example bone containing breast cancer metastases, collected *in situ* on the AFM. The MT differs from the surrounding tissues in colour in the conventional image (outlined by *dashed line* in f). The more accurate tumour region is identified from the green fluorescence protein (GFP) expressed by the cancer cells in MT (outlined by *dashed line* in g). The bone metaphysis region surrounding the MT was only accessed at distances greater than 200 µm from the fluorescent tumour edge (outlined by *dash-dotted lines* in f and g) in AFM measurements to avoid involving any cancer cells. (h) Conventional and (i) the corresponding *in situ* fluorescent images of a subcutaneous tumour (SCT) surface (outlined by *dashed lines*) immobilised on substrate. Cancer cells in SCT express GFP and can be identified from the fluorescent image (*arrows*). (h) Optical image of the MDA-MB-231^luc/GFP^ cells (*darker areas*) grown in 2D culture in a petri-dish.

Features associated with plastic deformation (including yield points, plateaus) were rarely observed from the *F*-*δ* curves. The *Δδ*-*t* curves from 93% of all measured positions were fitted well by the K-V model (*R*^2^ > 0.9). This demonstrates the MT in bone is viscoelastic and acts like a Kelvin-Voigt solid.

Histograms of *E*_*H-S*_, *E*_*K-V*_ and *η* of the MT are shown in Fig. 2 (note the logarithmic scale of the horizontal axes), all show non-normal distributions. The median values of *E*_*H-S*_, *E*_*K-V*_ and *η* are 5.2 Pa, 28 Pa and 17 Pa·s, while the mean values are 17 Pa, 65 Pa and 25 Pa·s, respectively(n=117 for *E*_*H-S*_; n=118 for *E*_*K-V*_ and *η*). These data indicate that overall, the MT is extremely compliant (i.e. has both low elastic modulus and low viscosity).

**Figure 2.**
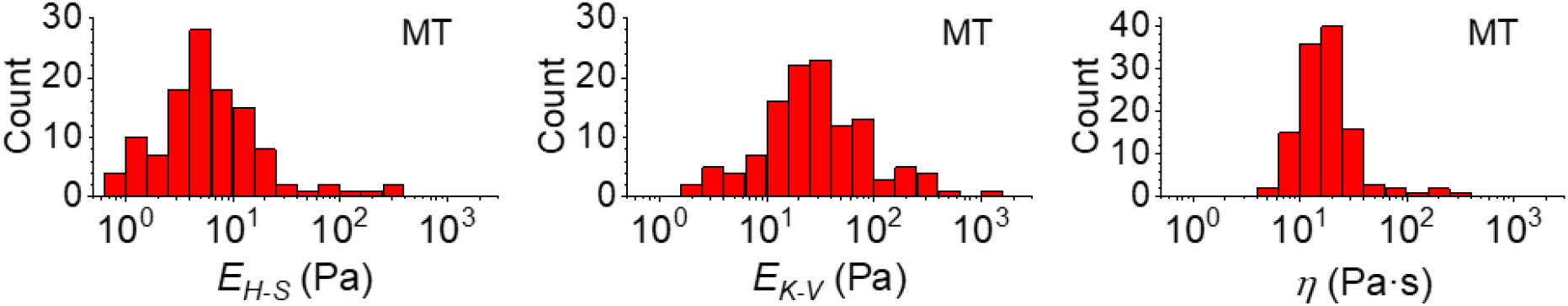
Histograms of the mechanical properties of the metastatic breast tumour in bone (MT) measured by AFM. The Young’s modulus *E*_*H-S*_ was calculated from the Hertz-Sneddon model fitting to the force-indentation (*F-δ*) curves measured at randomly selected positions within regions of interest. The Young’s modulus *E*_*K-V*_ and viscosity *η* were obtained from the Kelvin-Voigt model fits to the creep curves measured at the same positions as the *F-δ* curves. Each count of *E*_*H-S*_, *E*_*K-V*_ and *η* in the histograms is the mean value of individual fits to all force curves (≥ 3 repeats) taken at one position. Data were collected at n=126 positions from 19 bones from 16 tumour bearing mice. Results from low quality fittings (i.e. *R*^2^ < 0.9) have been discarded (~ 7% of all measurements).

The distributions of both *E*_*H-S*_ and *E*_*K-V*_ range over 2 orders of magnitude, while the width of *η* distribution is slightly narrower (approximately 83% of data points lying within 1 order of magnitude). This reveals mechanical heterogeneity across the whole of the MT. In contrast to findings from primary breast tumours^10^, the distributions of all three mechanical parameters lack any apparent second peaks. This is in agreement with the single peak of stiffness distribution determined from lung metastasis observed in the same study^10^. Furthermore, histological sections (Fig. 1d-e) from this bone metastases model show no focal regions of tumour fibrosis (collagen deposition), a common feature of primary human breast tumours that may contribute to the peak at higher stiffness observed/determined in the mechanical distribution of the primary tumour.

The creep time *τ* is defined as the time for the strain to decay to 1/*e* of its total change. We calculated the creep time by τ = 3*η*/*E*_*K*-*V*_ for each measured position. The resultant *τ* ranges between 0.3 and 13.7 s and the median value is 1.8 s. Acting like a Kelvin-Voigt solid, the MT is predominantly a viscous liquid at short time scales (*t* ≪ τ) and an elastic solid at long time scales.

Taken together, our findings demonstrate that AFM can be used to determine the mechanical properties of highly complex tissues like bone metastases, and reveal that metastatic breast tumours in bone have extremely low elastic modulus and viscosity.

### The microenvironment in which a tumour grows has significant impact on its mechanical properties

Over the past decades, the role of the multiple components of the microenvironment in tumour development and response to therapy has been increasingly recognised. As a result, ‘the hallmarks of cancer’ have been expanded to include interactions with the tumour microenvironment^22^. The environment in which the tumour grows is thus likely influencing its characteristics, including the mechanical properties^23^. Cancer researchers using different models (e.g. *in vitro* cultures of cancer cell lines, *in vivo* tumour cell growth) do not always use orthotopic transplantation for the tumour i.e. tumour growth in the normal site/tissue of origin. The mechanical contribution from the tumour microenvironment in cancer models implanted in a non-orthotopic site, compared to the appropriate host microenvironment, has not been well documented. We therefore used our AFM protocol to quantify the mechanical properties of breast tumours grown in different microenvironments, by comparing MDA-MB-231^luc/GFP^ cells growing as bone metastases (i.e. MT, measured at 126 positions from 19 bones), as non-orthotopic tumours (i.e. subcutaneous tumour established from MDA-MB-231^luc/GFP^ implantation as shown in Fig. 1h-i, measured at 209 positions from 8 tumours) or in 2D cultures (i.e. isolated MDA-MB-231^luc/GFP^ cells in a petri-dish as shown in Fig. 1j, measured on 95 cells from 7 petri-dishes).

*E*_*H-S*_, *E*_*K-V*_ and *η* measured on the MT, the subcutaneous tumour (SCT) and the MDA-MB-231^luc/GFP^ cells in 2D culture are represented in Fig. 3a, and demonstrate a highly significant difference between the three models (*p* < 0.001). The corresponding histograms are shown in Fig. 3b&S1. The median values of *E*_*H-S*_, *E*_*K-V*_ and *η* are (i) MT: 5.2 Pa, 28 Pa and 17 Pa·s, (ii) SCT: 11 Pa, 60 Pa and 26 Pa·s, (iii) MDA-MB-231^luc/GFP^ cells: 152 Pa, 559 Pa and 168 Pa·s. The mean values of *E*_*H-S*_, *E*_*K-V*_ and *η* are (i) MT: 17 Pa, 65 Pa and 25 Pa·s, (ii) SCT: 26 Pa, 145 Pa and 52 Pa·s, (iii) MDA-MB-231^luc/GFP^ cells: 204 Pa, 766 Pa and 205 Pa·s (n=117, 186 and 95 for *E*_*H-S*_ of MT, SCT and MDA-MB-231 cells; n=118, 198 and 93 for *E*_*K-V*_ and *η* of MT, SCT and MDA-MB-231 cells).

**Figure 3.**
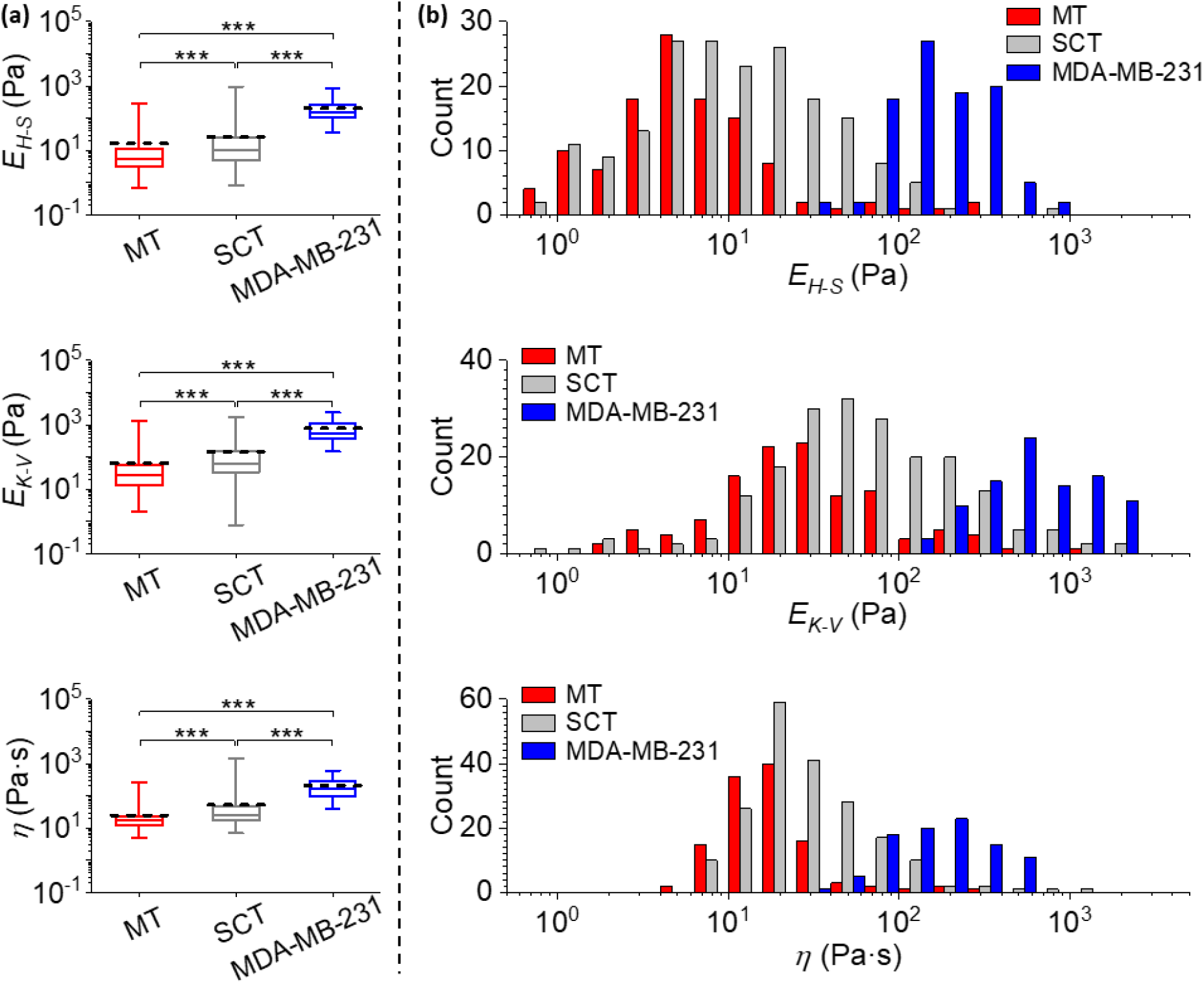
Comparisons of mechanical properties measured from different cancer models. (a) Statistical comparisons of the Young’s moduli *E*_*H-S*_, *E*_*K-V*_ and viscosity *η* of metastatic breast tumour in bone (MT, data are identical to those in Fig. 2), subcutaneous tumour (SCT) and MDA-MB-231^luc/GFP^ cells grown in 2D cultures in petri-dishes (***: *p* < 0.001). Data were analysed using the same method as in Fig. 2, and collected from biological and technical repeats (MT: n=126 positions from 19 bones of 16 mice; SCT: n=209 positions from 8 tumours established in 6 mice; MDA-MB-231: n=95 cells cultured in 7 petri-dishes). The central box spans the lower to upper quartile of the data. The solid line inside the box represents the median and whiskers represent the lower and upper extremes. The mean values are indicated by dashed lines. Note the logarithmic scale of the y axes. Results from low quality fittings (i.e. *R*^2^ < 0.9) were discarded (~ 7%, 11% and none of all measurements for *E*_*H-S*_ of MT, SCT and MDA-MB-231 cells; ~ 7%, 5% and 2% of all measurements for *E*_*K-V*_ and *η* of MT, SCT and MDA-MB-231 cells). (b) Histograms of the *E*_*H-S*_, *E*_*K-V*_ and *η* of the MT, SCT and MDA-MB-231^luc/GFP^ cells, correspond to data in (a). Bars in each histogram were narrowed down and shifted (i.e. a group of 3 subset bars, in order of MT, SCT and MDA-MB-231, has the same bin size in reality that equals the total width of 3 bars in the x-axis scale) to avoid stacking of columns. Note the logarithmic scale of the x axes. Individual histograms are shown in the Supporting Material.

It should be noted that all measured mechanical properties of MDA-MB-231^luc/GFP^ cells grown in 2D culture have more than an order of magnitude higher peak values in the histograms, when compared to those of 3D explanted tumours (i.e. both MT and SCT). In contrast to a previous study demonstrating that isolated tumour epithelial cells are softer than the tumour epithelium measured *in situ*^24^, our results indicate cancer cells are significantly more compliant in a bone tissue environment when compared to isolated 2D cultured cells. The indentation depth of our *in vitro* cultured cancer cells (< 640 nm for cells with median *E*_*H-S*_ or above) is significantly smaller than the cell dimensions. Therefore, we estimate the influence of the hard substrate on the AFM indentation measurements is negligible, and the resultant higher elastic moduli of MDA-MB-231^luc/GFP^ cells compared to the 3D tumours, is a consequence of the 2D cell culture (i.e. cells adhere to a hard surface and consequently under tension) rather than an artefact arising from the petri-dish substrate. Meanwhile, the width between upper and lower quartiles of the elastic moduli is narrower in isolated cancer cells, but the width of viscosity is similar. This suggests that in 3D tumours other components (e.g. extracellular matrix) contribute to an increased heterogeneity of the elasticity.

It is interesting that the shapes of *E*_*H-S*_, *E*_*K-V*_ and *η* distributions are almost identical between SCT and MT. However, where MT has lower mean and median values of the three parameters quantified, this was statistically significant (Fig. 3a) when compared to SCT. These data indicate that the metastatic niche in bone significantly enhances the tumour compliance, even though it does not significantly affect the degree of tumour heterogeneity.

Immunofluorescent staining of the extracellular matrices (ECM) commonly associated with breast cancer metastases in bone (i.e. Collagen I, Collagen IV and Laminin) was used to identify ECM proteins on tissues we had characterised using AFM (see Supporting Material, Fig. S2). This evaluation aimed to determine if the differential mechanical measurements were associated with different ECM deposition in the respective tissues. There was an abundance of all three extracellular components in the SCT when compared to the MT, though statistical significance was only identified for Collagen I and Laminin. Therefore, increasing ECM deposition is positively correlated with the measured *E*_*H-S*_, *E*_*K-V*_ and *η* of tumour, which is in agreement with findings from previous studies^23, 24, 25^.

Overall these data demonstrate that metastatic breast tumours in bone are significantly more compliant than both 3D subcutaneous breast tumours and 2D breast cancer cells *in vitro*, supporting the notion that the microenvironment in which the tumour grows, impacts on the resultant mechanical properties.

### Does the tumour alter the mechanical properties of the surrounding bone microenvironment?

Breast tumour growth in bone is associated with increased bone resorption, resulting in lytic lesions and weakening of the bone, termed cancer-induced bone disease (CIBD)^13^. Although the detrimental effects of CIBD on calcified bone are widely studied, there is limited knowledge on how the presence of a tumour mechanically influences the surrounding microenvironment, including not only calcified bone but also multiple types of cells and extracellular matrices. Our mouse model of breast cancer bone metastasis also allowed mechanical characterisation of the relatively intact microenvironment surrounding the MT.

We focused on the bone metaphysis region (bone surrounding tumour, measured at 107 random positions from 19 bones). To assess the normal tissues without any cancer cells, this region was characterised at distances greater than 200 µm from the fluorescent tumour edge (Fig. 1f-g, dash-dotted region). In addition, the same mechanical measurements were made in the bone metaphysis from non-tumour bearing mice acting as a negative control (bone w/o tumour, measured at 192 random positions from 22 bones), to reveal the mechanical impact of tumours on the bone metastatic niche. All measurements were collected at randomly selected positions within the bone metaphysis region, including both bone tissues and bone marrow.

Statistical comparisons of *E*_*H-S*_, *E*_*K-V*_ and *η* measured on the MT, bone surrounding tumour and bone w/o tumour are shown in Fig. 4a. The corresponding histograms are shown in Fig. 4b&S1. The median values of *E*_*H-S*_, *E*_*K-V*_ and *η* are (i) MT: 5.2 Pa, 28 Pa and 17 Pa·s, (ii) bone surrounding tumour: 17 Pa, 84 Pa and 34 Pa·s, (iii) bone w/o tumour: 25 Pa, 100 Pa and 41 Pa·s. The mean values of *E*_*H-S*_, *E*_*K-V*_ and *η* are (i) MT: 17 Pa, 65 Pa and 25 Pa·s, (ii) bone surrounding tumour: 75 Pa, 140 Pa and 79 Pa·s, (iii) bone w/o tumour: 165 Pa, 452 Pa and 108 Pa·s (n=117, 86 and 154 for *E*_*H-S*_ of MT, bone surrounding tumour and bone w/o tumour; n=118, 96 and 143 for *E*_*K-V*_ and *η* of MT, bone surrounding tumour and bone w/o tumour).

**Figure 4.**
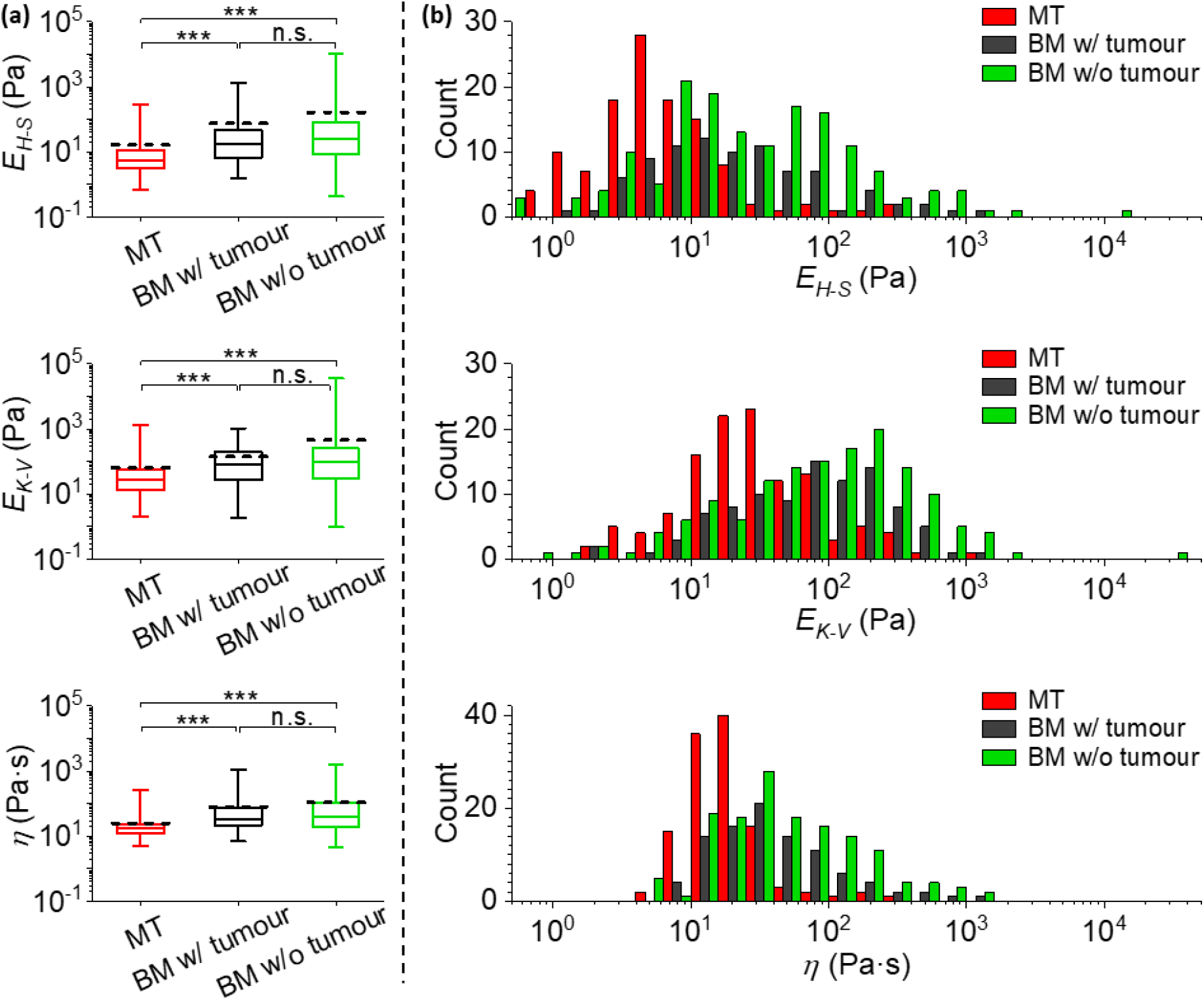
The mechanical comparisons between the metastatic tumour and the surrounding tissue microenvironment. (a) Statistical comparisons of the Young’s moduli *E*_*H-S*_, *E*_*K-V*_ and viscosity *η* of metastatic breast tumour in bone (MT, data are identical to those in Fig. 2), bone metaphysis surrounding tumour (BM w/ tumour, as in Fig. 1f-g) and bone metaphysis from tumour-free mice (BM w/o tumour) (***: *p* < 0.001; n.s.: no significance). Data were analysed using the same method as in Fig. 2, and collected from biological and technical repeats (MT: n=126 positions from 19 bones of 16 tumour bearing mice; BM w/ tumour: n=107 positions from the same 19 bones as used for MT; BM w/o tumour: n=192 positions from 22 bones of 11 non-tumour bearing mice). The central box spans the lower to upper quartile of the data. The solid line inside the box represents the median and whiskers represent the lower and upper extremes. The mean values are indicated by dashed lines. Note the logarithmic scale of the y axes. Results from low quality fittings (i.e. *R*^2^ < 0.9) were discarded (~ 7%, 20% and 20% of all measurements for *E*_*H-S*_ of MT, bone surrounding tumour and bone w/o tumour; ~ 7%, 10% and 25% of all measurements for *E*_*K-V*_ and *η* of MT, bone surrounding tumour and bone w/o tumour). (b) Histograms of the *E*_*H-S*_, *E*_*K-V*_ and *η* of the MT, BM w/ tumour and BM w/o tumour, correspond to data in (a). Bars in each histogram were narrowed down and shifted (i.e. a group of 3 subset bars, in order of MT, BM w/ tumour and BM w/o tumour, has the same bin size in reality that equals the total width of 3 bars in the x-axis scale) to avoid stacking of columns. Note the logarithmic scale of the x axes. Individual histograms are shown in the Supporting Material.

All the tissues measured demonstrate a high level of compliance, however the MT is significantly more compliant than both bone surrounding tumour and bone w/o tumour (*p* < 0.001). The widths between upper and lower quartiles of all MT mechanical parameters are narrower than those of the surrounding bone metaphysis. This is consistent with previous published data using primary tumours, where regions rich in cancer cells were more compliant and less heterogeneous than the surrounding normal tissues^10, 26^.

As previously described, we investigated whether differences in the biomechanical properties between MT and the surrounding bone environment were due to differences in extracellular matrix composition. Immunofluorescent staining of the extracellular components was quantified and compared between MT and the non-tumour sites in the tumour bearing bone metaphysis (see Supporting Material, Fig. S2). No significant difference was observed in the amount of Collagen I, Collagen IV and Laminin between the different samples. This reveals that the mechanical distinction between MT and its tissue environment is unlikely to result from the ECM proteins evaluated in this study. However, different cell types and other extracellular components may still contribute to the differences observed and should be explored in future studies.

Interestingly, the shapes of *E*_*H-S*_, *E*_*K-V*_ and *η* distributions are relatively similar between bone surrounding tumour and bone w/o tumour. No statistically significant differences are observed from the overall distributions, though the mean values of *E*_*H-S*_ and *E*_*K-V*_ are greater for the bone w/o tumour compared to the bone surrounding tumour (most likely resulting from the low number of discrete data points at the higher extreme). We find no evidence that the presence of a tumour in bone significantly influences the mechanical properties of the remote (> 200 µm from tumour) microenvironment, even though the impact within a shorter range remains unknown.

## DISCUSSION

We have developed a robust AFM based method^21^ and successfully used this to quantify the mechanical properties of relatively intact breast cancer bone metastases from an *in vivo* mouse model^15, 20^. Our results show that the MT possesses an extremely low elastic modulus (down to a few Pa) and viscosity (down to a few Pa?s), even when compared to the highly compliant healthy murine bone soft tissue^21^. The resultant elastic modulus is several orders of magnitude lower than that measured at the macroscopic scale^8^, and is likely more relevant to cellular processes.

Although widely used to study molecular mechanisms of cancer development, *in vitro* cultures of cancer cell lines have often been criticised as being over-simplified cancer models. Consisting of a single cell-type grown in 2D, these cell models do not represent the complex, multicellular 3D *in vivo* tumour environment. It is thus essential to quantify any differences between the mechanical properties of 3D *in vivo*/*ex vivo* tumours and the relevant 2D *in vitro* cell model, in order to improve research models and hence increase our understanding of the mechanical processes involved in tumour progression. The challenge is exacerbated by the wide variation in Young’s moduli determined from the same *in vitro* cell lines published by different laboratories^5, 27, 28, 29^, making a direct comparison of the properties measured in the current study difficult. Due to likely variations in the same cell-line (e.g. following genetic manipulation and clonal selection), we quantified the mechanical properties of isolated MDA-MB-231^luc/GFP^ cells (the same cell line used to establish the bone metastases and the subcutaneous tumours in mice) in a petri-dish under the same experimental conditions as used for the tumours (e.g. same temperature, same AFM settings). The median *E*_*H-S*_, *E*_*K-V*_ and *η* of MDA-MB-231^luc/GFP^ cells is increased when compared to the MT by 29, 20 and 10 times respectively. This demonstrates the cancer cells are significantly stiffer in a 2D *in vitro* culture than when the same cell line is grown in 3D at a metastatic site. This should be taken into account in future studies using 2D *in vitro* cancer models and highlights the need for methods to perform mechanical analysis in complex tissues.

Moreover, we characterised both orthotopic (i.e. MT) and non-orthotopic (i.e. SCT) tumour models to determine any differential mechanical properties between different implantation sites and whether an appropriate breast metastatic niche (e.g. bone) possesses additional mechanical cues. Although the bone niche does not affect the degree of tumour heterogeneity, MT has a significantly reduced elastic modulus and viscosity when compared to SCT. This can be explained by several potential mechanisms. The first possibility is that cancer cells can significantly alter their mechanics, including the extracellular components, when growing in different environments. This is supported by published reports demonstrating that orthotopic breast cancer xenografts have greater elasticity and viscosity compared to tumours associated with the nervous system^23^. Secondly, metastases are potentially associated with stiffness selection. Previous studies using both 2D *in vitro* cell models^30^ and tumour tissue models^25, 31^ suggest that more compliant cancer cells and primary tumours are associated with enhanced tumour progression and extensive metastases. Therefore, it is possible that the MT in bone develops from a more compliant subpopulation present within the injected cancer cells. This potential mechanism may not work in isolation but be associated with other mechanisms, because the minimum elastic moduli and viscosity of isolated MDA-MB-231^luc/GFP^ cells measured in this study is higher than the majority (i.e. upper quartile) of those of MT. Finally, yet importantly, the surrounding tissue microenvironment in the bone niche may also alter the mechanical properties of cancer cells that have reached bone, via physical and/or biochemical interactions.

By comparing the mechanical properties of MT and its surrounding bone microenvironment, we conclude that MT outgrowth favours tissue microenvironments that are less compliant (i.e. stiffer and more viscous) and more mechanically heterogeneous than tumour tissues. This offers a clear benchmark for designing more rationalised *in vitro* cancer research models and designing mechanical interventions as anti-cancer drugs/treatment, as suggested in a recent study on metastatic colorectal cancer to the liver^32^. In addition, the mechanical properties of the bone surrounding tumour (> 200 µm away from the tumour margin) show no significant difference compared to the bone w/o tumour. This implies that MT does not mechanically alter the tissue microenvironment at this distance, but we cannot exclude that the microenvironment at the tumour-bone interface may be affected. These findings are in agreement with those of a previous study using the same *in vivo* model, where we identified significant changes in bone cell numbers (osteoblasts and osteoclasts) only in the areas of bone that were in direct contact with the tumour^33^. It will therefore be important to map the local mechanical architecture at a range shorter than ~200 µm from the tumour/bone tissue interface in future studies.

In this work, we have combined *in vivo* models of breast cancer spread to bone with AFM and shown that this approach provides valuable information about the mechanical properties of the breast cancer metastases together with its surrounding bone microenvironment as nearly intact complex tissues. By comparing to non-orthotopic tumour, isolated cancer cells and bone w/o tumour, we have established clear benchmarks of the mechanical relation between the metastases and its microenvironment, which are fundamental for designs of future studies. We expect that this methodology can be used to increase our understanding of the mechanism of cancer metastasis, as well as how this may be targeted through disruption of the mechanical interactions with the bone microenvironment.

## METHODS

### Animals

All experiments involving animals were approved by the University of Sheffield Project Applications and Amendments (Ethics) Committee and conducted in accordance with UK Home Office Regulations (PPL70/8964 to N.J. Brown). Female BALB/c Nude [Foxn1-Crl] immunodeficient mice (Charles River, UK) (n=27 in total) were housed in a controlled environment with 12 hours light/dark cycle, at 22 °C. Mice had access to food and water *ad libitum*.

### Sample preparation for different cancer models

Green Fluorescent Protein (GFP) expressing triple negative breast cancer cells (MDA-MB-231^luc^/^GFP^) were cultured in RPMI-1640 GlutaMAX™ medium (Thermo Fisher Scientific, Waltham, MA, USA) supplemented with 10% FCS, maintained at 37 °C and 5% CO_2_. Cells harvested from the subculture were seeded into petri-dishes (TPP, Switzerland) one day prior to mechanical measurements and maintained in the incubator. The medium was changed immediately prior to mechanical measurements to remove any dead cells.

Subcutaneous tumours (SCT) were established by injecting 1×10^6^ MDA-MB-231^luc/GFP^ into the hind flank. Mice were culled 4 to 5 weeks post-injection when palpable tumours were detected for analysis.

Experimental bone metastatic tumour (MT) were established in mice (6 weeks old) by injecting 5×10^4^ MDA-MB-231^luc/GFP^ cells into the left cardiac ventricle, as described previously^15, 20^. The growth of bone metastases was monitored twice weekly post-injection by non-invasive bioluminescent imaging using an IVIS Lumina II imaging system (Fig. 1a). Mice were culled for analysis when a positive luciferase signal was monitored in the hind limbs (within 3 to 4 weeks post-injection). The size of tumours in bone was variable, but a mature tumour was observed in most cases if a strong bioluminescent signal was obtained on bones from which the attached soft tissues had been removed (Fig. 1b). Bones from the same strain of non-tumour bearing mice (6 to 8 week old) were used as the negative control (bones w/o tumour).

Dissected subcutaneous tumours and hind limbs (both femurs and tibias) were placed in phosphate buffered saline (PBS) (Lonza, US) at 4 °C. Both types of specimens were split using a razor blade and immobilised in a petri-dish using a two-component dental impression putty (Provil Novo Light, Kulzer, UK), immediately before mechanical measurements. Care was taken to maintain hydration of the exposed sample surface during the entire process.

### Force-indentation and creep analyses by AFM

Both elastic and viscoelastic properties of different cancer models were determined from force-indentation (*F*-*δ*) and creep curves obtained by atomic force microscopy (AFM), using the method described previously^21^. AFM measurements were performed on a Nanowizard III system (JPK, Germany) with an extended z piezo range (100 µm), combined with an inverted optical microscope (Eclipse Ti, Nikon, Japan) and a custom built top view optical microscope (broadband and fluorescent imaging, excitation: 445/45 nm, emission: 525/39 nm). Details of the long-range Z scanner and top view optics are provided in Methods of Supporting Material. Rectangular cantilevers (MLCT-Bio-DC, cantilever B with nominal spring constant of 0.02 N/m) (Bruker, USA) with a 25 µm diameter polystyrene microsphere (Sigma Aldrich, USA) glued to the free end were used in all AFM measurements, after passivation with 10 mg/mL bovine serum albumin (BSA), (Sigma Aldrich, USA). The spring constant and deflection sensitivity of each cantilever was calibrated prior to each measurement^34^.

Measurements were performed at 36 to 37 °C. SCTs and bones with metastases remained in PBS and MDA-MB-231^luc/GFP^ cells in culture medium throughout the measurements. AFM data on a single sample (i.e. one piece of SCT or bone, or one petri-dish of MDA-MB-231^luc/GFP^ cells) could typically be collected in 2 to 3 h. All tissue measurements were completed no longer than 12 hours post-cull.

*In situ* optical images (Fig. 1f-j) aided targeting the area of analysis. For tissues, *F*-*δ* curves were acquired at randomly selected positions within different regions of interest, including whole metaphysis of non-tumour bearing bone (Fig. 1c), MT (Fig. 1f-g, dashed region), bone metaphysis surrounding the tumour at a distance greater than 200 µm (Fig. 1f-g, dash-dotted region) and SCT (Fig. 1h-i). For MDA-MB-231^luc/GFP^ cells in petri-dishes, *F*-*δ* curves were acquired on top of the nuclei (Fig. 1j). The trigger force was 0.5 nN and the approach speed 5 µm/s. Subsequently, curves with 3 s dwell under constant force (0.5 nN), *i*.*e*. creep curves, were also acquired from the same position. For both *F*-*δ* and creep curves, a minimum of 3 measurements were taken at each location.

Raw data were exported as .txt format using JPK Data Processing and imported into customised algorithms in MATLAB for all subsequent analyses. As described previously^21^, *F*-*δ* curves were fitted to a Hertz-Sneddon (H-S) model^35^ to acquire the Young’s modulus (*E*_*H-S*_) assuming a virtual contact point, and creep curves was fitted to a Kelvin-Voigt (K-V) model^36^to extract the Young’s modulus (*E*_*K-V*_) and viscosity (*η*). Results were obtained from the mean value of repeated measurements at each position and those with low fit quality (*R*^2^ < 0.9) were discarded.

### Statistics

Data of MT and its surrounding bone metaphysis were collected at 126 and 107 positions respectively, from 19 bones resected from 16 tumour bearing mice. Data from bones w/o tumour were acquired at 192 positions from 22 bones of 11 non-tumour bearing mice. Data from SCT were collected at 209 positions from 8 tumours established in 6 mice and data for MDA-MB-231^luc/GFP^ cells from 95 cells in 7 petri-dishes. The mechanical properties of each measured position/cell were determined from the mean value of the analysed results of repeated force curves taken at each location (≥ 3 repeats). The analysed results of each force curve with low fitting quality (i.e. *R*^2^ < 0.9) were discarded. The N numbers (i.e. number of measured positions/cells) of data obtained from H-S model (i.e. *E*_*H-S*_) are: (i) MT: 117, (ii) SCT: 186, (iii) MDA-MB-231: 95, (iv) bone surrounding tumour: 86, (v) bone w/o tumour: 154. The N numbers of data obtained from K-V model (i.e. *E*_*K-V*_ and *η*) are: (i) MT: 118, (ii) SCT: 198, (iii) MDA-MB-231: 93, (iv) bone surrounding tumour: 96, (v) bone w/o tumour: 143. Statistical analyses were performed using OriginPro software. A normality test was applied to all distributions prior to any further analysis. Data were analysed by the Kruskal-Wallis test for comparison between different groups. A statistically significant difference was defined as *p* < 0.05.

## AUTHOR CONTRIBUTIONS

XC designed the study, performed the experiments, analysed and interpreted the data, and prepared the manuscript. RH prepared the biological samples, interpreted the data and contributed to manuscript preparation. NM designed and constructed the top-view optics and extended z-stage, interpreted the data and contributed to manuscript preparation. RJH improved the theoretical model, interpreted the data and contributed to manuscript preparation. IH, NJB and JKH designed the study, interpreted the data, contributed to manuscript preparation and directed the project.

## ACKNOWLEDGEMENTS

This research was supported by Cancer Research UK and the Engineering and Physical Sciences Research Council (Grant Number: A21082). We thank Prof. Keith Hunter, Professor. Ashley Cadby and Miss Natasha Cowley (University of Sheffield) for fruitful discussions. We also thank Dr Heiko Haschke (JPK Instruments) for assistance with integrating the long-range Z scanner and Simon Dixon and Mark Lister (Physics Workshop, University of Sheffield) for fabrication of bespoke components for the long-range Z scanner and top-view optics.

## COMPETING INTERESTS

We declare no competing interests relevant to this work.

## Notes

### Competing Interest Statement

The authors have declared no competing interest.

