## Supporting Material for "Atomic force microscopy reveals the mechanical properties of breast cancer bone metastases"

### SUPPORTING METHODS

#### Immunofluorescent imaging and image analysis

Bone tissue was fixed and maintained in 4% paraformaldehyde (PFA) at 4ºC, following analysis by AFM. Fixed bones were decalcified in 0.5 M EDTA pH 7.5 overnight at 4ºC. Decalcified bones were equilibrated in cryoprotectant (20% sucrose and 2% PVP in PBS) overnight at 4ºC before being embedded in gelatin (8% porcine gelatin, 20% sucrose and 2% PVP in PBS) at 60ºC and then stored at -80ºC. Gelatin-embedded bones were cut into 30 µm thick cryo-sections and immunofluorescent staining was performed on three non-sequential sections. For immunofluorescent labelling, cryo-sections were defrosted, rehydrated in PBS, permeabilised in 0.3% Triton-X100 for 20 minutes at ambient temperature, washed in PBS and treated with a streptavidin/biotin blocking kit (Vector Labs, CA, USA). Tissue sections were then incubated with rabbit polyclonal primary antibodies against Collagen I, Collagen IV and Laminin (AbCam, Cambridge, UK) at a concentration of 10 µg/mL for 2 hours at ambient temperature. Tissue sections were then incubated with biotinylated goat anti-rabbit secondary (5 µg/mL) for 1 hour at ambient temperature, washed with PBS and incubated with Alexa Fluor 555-conjugated streptavidin (10 µg/mL) for 30 minutes (Thermo Fisher Scientific, MA, USA). Labelled sections were stained with DAPI, washed in PBS and mounted in ProLong Diamond antifade (Thermo Fisher Scientific, MA, USA).

Cryo-sections from subcutaneous tumours were defrosted and fixed in 4% PFA solution at ambient temperature for 10 minutes, permeabilised with 0.1% Triton X-100 for 5 mins, then washed in PBS. Subcutaneous tumour cryo-sections were then immuno-fluorescently labelled as described above.

Images of stained tissue sections were acquired using a Zeiss LSM 980 (Zeiss Group, Germany) confocal system. Sections from more than 3 tissues for each sample type were imaged. For bone sections, imaging was focused on the metaphysis region.

Images were analysed using ImageJ (National Institute of Health, US) to quantify the fluorescently labelled extracellular components. Statistical analyses were performed using Prism software (v.8.0, GraphPad Software, La Jolla, CA, USA). All data are presented as sample means ± SEM and were analysed using one-way ANOVA. A statistically significant difference was defined as $p < 0.05$.

### Top view optics combined with AFM

The opaque samples studied here required top view optics capable of both conventional and fluorescence imaging with a wide field of view. A bespoke mount was used to secure a simple optical microscope onto the condenser stage of a Nikon Eclipse Ti. 25.4 mm diameter cemented achromatic doublets (Thorlabs) with focal lengths of 75 mm and 200 mm were used for the objective and tube lenses, respectively, to give a 2.67x magnification and a field of view of approximately 1.8 mm x 1.35 mm with the camera used (Imagingsource DFK31BF03, 1/3” sensor). The ~67 mm working distance of the objective lens was sufficient to allow imaging through the AFM head. A filter holder was placed in the infinity space between the objective and tube lenses for the emission filter. To minimise contrast loss from scattered light from the optical components in the AFM head, oblique illumination was used, provided by a CooLED pE300 light source and delivered via a liquid light guide whose output was passed through the excitation filter, then weakly focused and steered onto the sample surface through the gap between the AFM head and stage using a kinematic tip/tilt mount. A photograph of the top view optics and AFM setup is shown in figure S3.

### AFM long Z scanner

Due to the large topography and extremely compliant nature of the samples, the 15 μm range of the Z piezo in the AFM head was not sufficient. In our setup, the Z piezo in the AFM head was disabled, and the sample was mounted on Z piezo stage with 100 μm of range (P-621, Physik Instrumente), which was controlled by the low voltage Z signal from the AFM passed through a high voltage amplifier (PI E500 with E505 HVA and E509 signal conditioner/piezo servo module, Physik Instrumente). Closed loop control was implemented in the E509, and the sensor monitor signal was passed to the AFM controller through the signal access module.

The long Z stage was calibrated by scanning 3 separate silicon calibration gratings in tapping mode, using the Z piezo in the AFM head and then the long Z stage immediately afterwards. A calibration factor to fix the calibration of the long Z stage such that the height sensor signal from the long Z stage matched that from the (factory calibrated) height sensor in the AFM head was calculated from height histograms of these measurements and set in the software. The measurement of the grating with the Z piezo within the AFM head agreed with the height specifications of the gratings within the manufacturer’s specified tolerances, and measurements after calibration showed agreement to within better than 0.5% between the calibrated long Z stage and the Z piezo in the AFM head.

A bespoke aluminium mounting plate was made to fit onto the chassis of the Nikon Eclipse Ti microscope in place of the standard AFM sample stage. A micrometer controlled XY stage (XYT1/M, Thorlabs) was mounted on this, to allow positioning of the sample relative to the AFM head. The long Z stage was mounted on top of this. On top of the long Z stage, a bespoke acrylic plate was used to stop any ingress of buffer into the stage from accidental spillages or leaks. A thin-film resistive heating element encapsulated in Kapton (KHA, 1.5 inch diameter, Omega Engineering) was glued to the top surface of acrylic plate with epoxy resin, and a bespoke copper petri-dish holder was glued to the top surface of the heater with thermal epoxy, to transfer heat to the sample in a petri dish. A Pt100 platinum resistance sensor, encapsulated in either glass (1PT100GO1020HG Omega Engineering) or alumina (HEL-705-T-0-12-00, Honeywell) was placed in the petri dish and held in place by a magnet glued to the sensor (glass encapsulated) or a small arm held in place by a magnet outside the dish (alumina encapsulated) and immersed in buffer. A commercial temperature controller (HTHS, JPK, which used a Eurotherm 2216e controller) was re-tuned to control the system, and found to maintain the temperature to within +/- 0.5 degrees of the set temperature by independent measurement with a type K thermocouple and DAQ card. A photograph of the constructed stage is shown in figure S4.

### SUPPORTING FIGURES

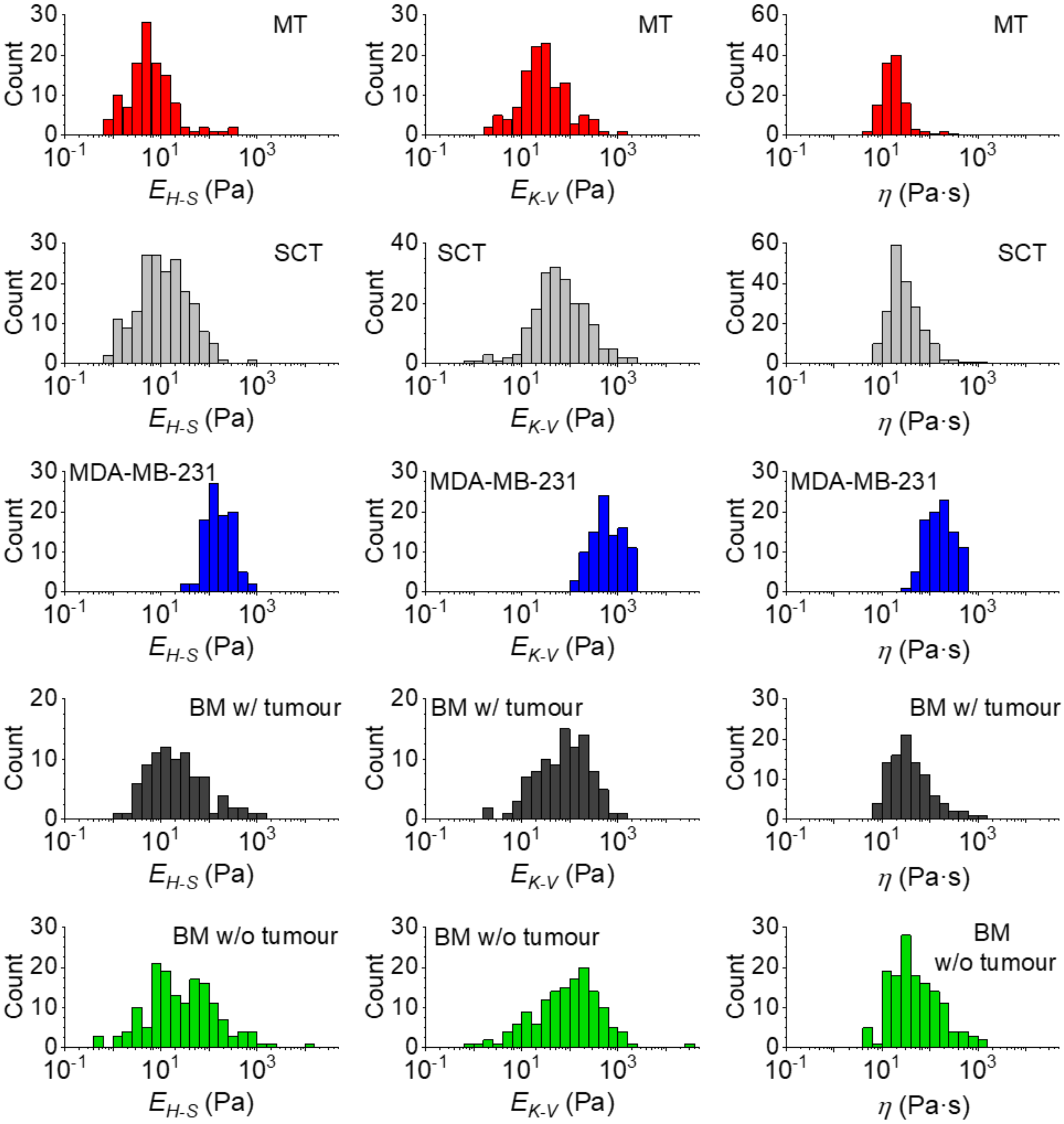

**Figure S1.** Histograms of the Young's moduli $E_{H\text{-}S}$, $E_{K\text{-}V}$ and viscosity $\eta$ of the metastatic tumour in bone (MT), subcutaneous tumour (SCT), MDA-MB-231$^{luc/GFP}$ cells grown in 2D culture in petri-dishes, the bone metaphysis surrounding tumour (BM w/ tumour) and bone metaphysis from tumour-free mice (BM w/o tumour). The data shown here are identical to those in Fig. 2 to 4.

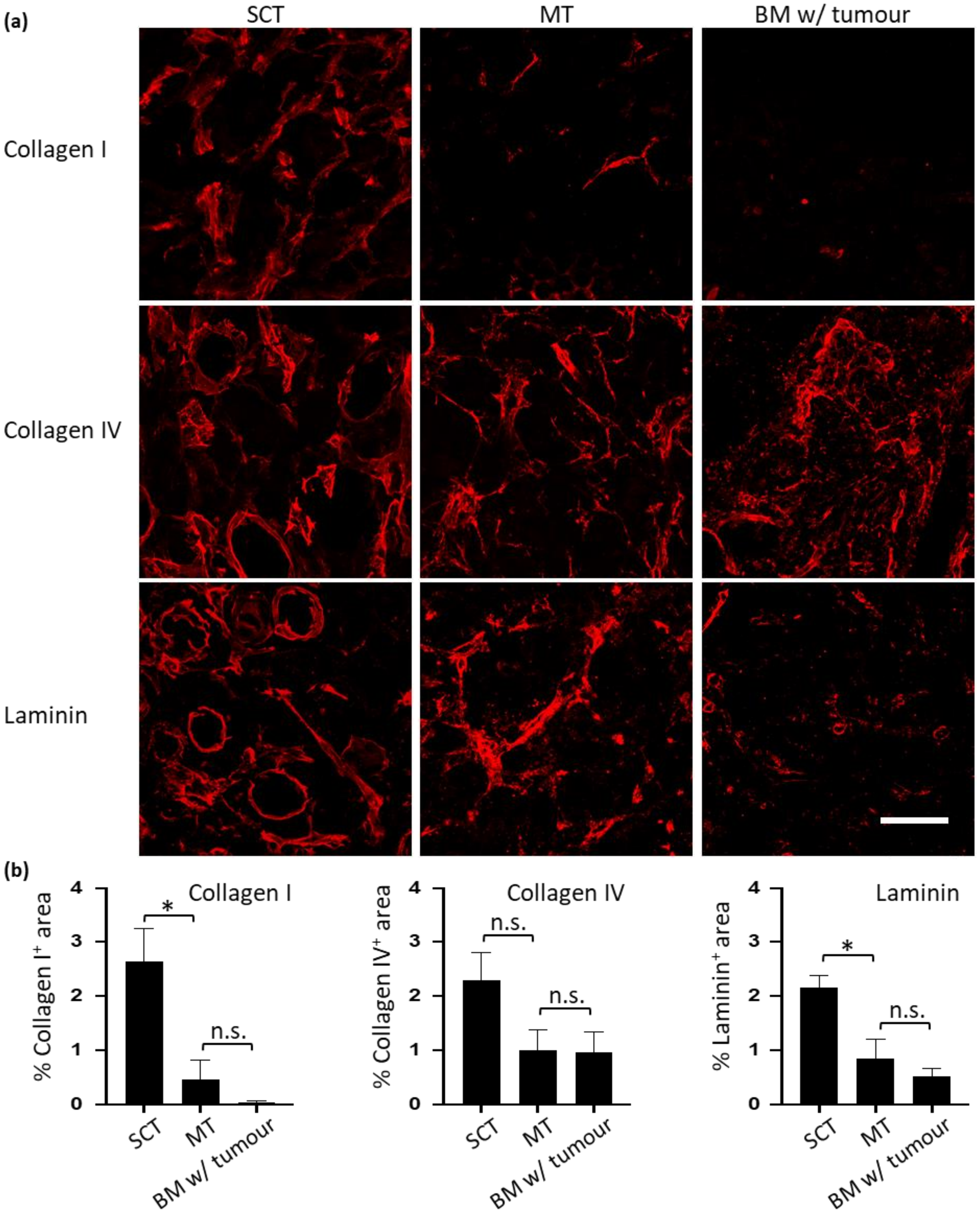

**Figure S2.** Immunofluorescent analysis of the extracellular matrices (ECM) potentially involved in cancer metastases. (a) Illustrative immunofluorescent images of Collagen I, Collagen IV and Laminin (all in *red*) obtained from the subcutaneous tumour (SCT), the metastatic tumour (MT) and the surrounding bone microenvironment (BM w/ tumour) (scale bar = 100 µm). (b) Quantification of different ECM components (**: $p < 0.01$; *: $p < 0.05$; n.s.: no significance; $n \geq 3$ per sample type).

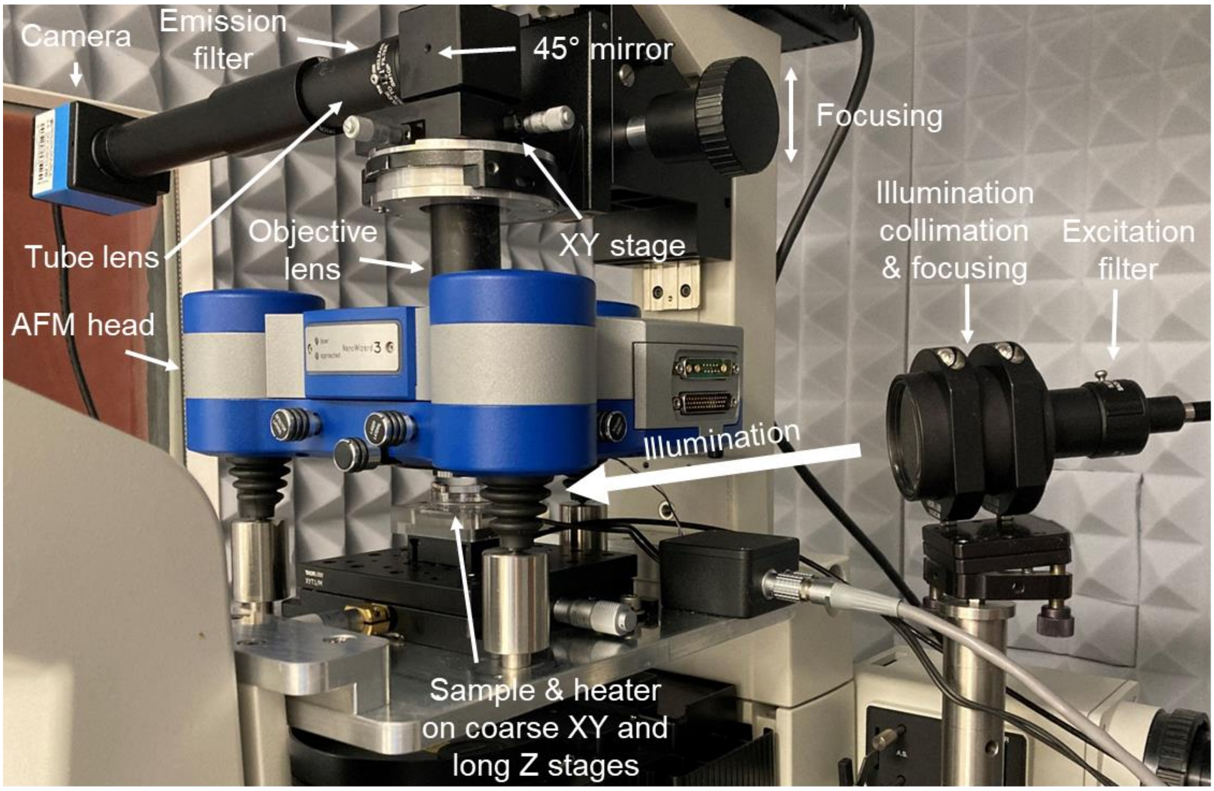

**Figure S3.** Annotated photograph of the AFM system with custom top view optics and 100 µm range Z piezo. The cables that connect the AFM head to the controller have been removed for clarity.

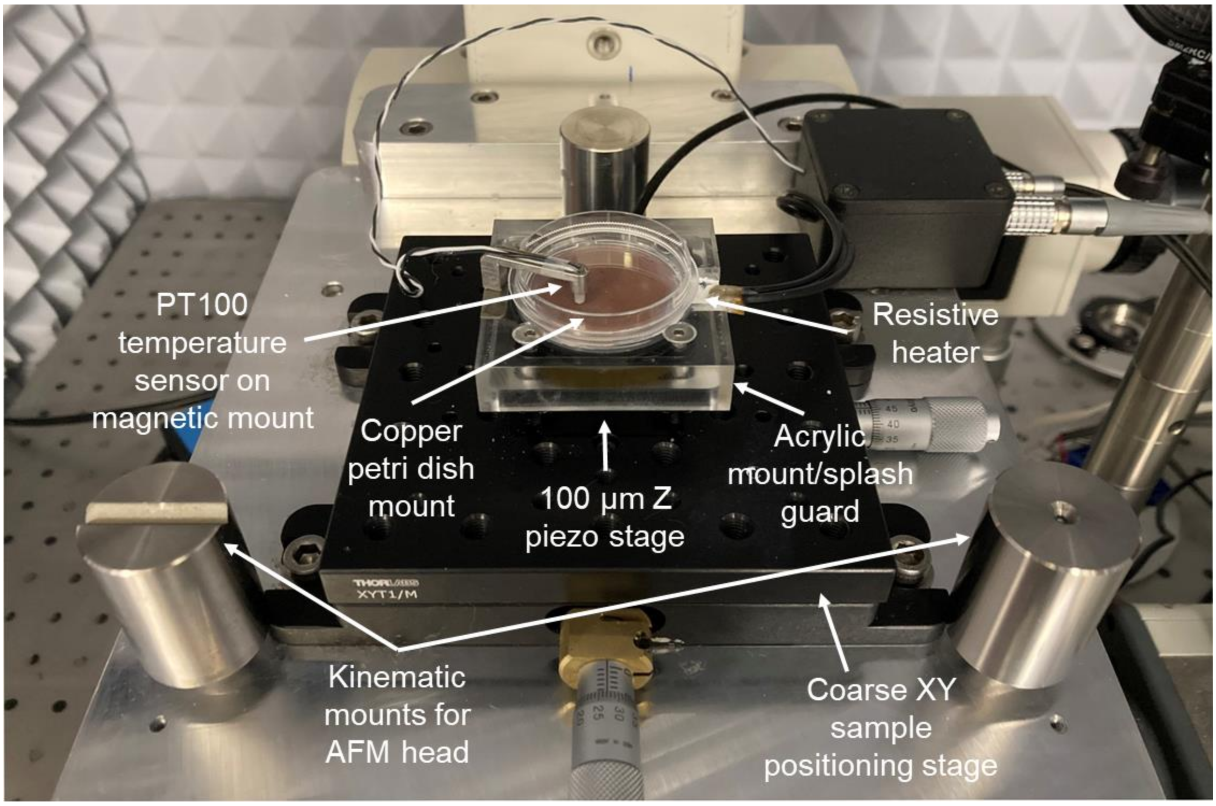

**Figure S4.** Annotated photograph of the long range Z stage, petri dish heater and coarse sample positioning stage. The AFM head has been removed for clarity. The long range z scanner is largely obscured by the acrylic mount/splash guard, and the resistive heater is obscured under the copper petri dish mount.
